# *M. leprae* infects human keratinocytes via the interaction of laminin-5 with α-dystroglycan, integrin-β1, or -β4

**DOI:** 10.1101/591313

**Authors:** Song-Hyo Jin, Se-Kon Kim, Seong-Beom Lee

## Abstract

Although *Mycobacterium leprae* (*M. leprae*) is usually found in macrophages and nerves of the dermis of patients with multibacillary leprosy, it is also present in all layers of the epidermis, basal, suprabasal, prickle cells, and keratin layers. However, the mechanism by which *M.leprae* invades the dermis remains unknown, whereas the underlying mechanism by which *M.leprae* invades peripheral nerves, especially Schwann cells, is well defined. *M. leprae* binds to the α-dystroglycan (DG) of Schwann cells via the interaction of α-DG and laminin (LN)-α2 in the basal lamina, thus permitting it to become attached to and invade peripheral nerves. In the current study, we investigated the issue of how *M.leprae* infects keratinocytes. LN-5 is the predominant form of laminin in the epidermis and allows the epidermis to be stably attached to the dermis via its interaction with α/β-DG as well as integrins that are produced by keratinocytes. We therefore focused on the role of LN-5 in when *M. leprae* invades keratinocytes. Our results show that *M.leprae* preferentially binds to LN-5-coated slides and this binding to LN-5 enhances its binding to human epidermal keratinocytes, neonatal (HEKn). The findings also show that pre-treatment with an antibody against α-DG, integrin-β1, or -β4 inhibited the binding of LN-5-coated *M.leprae* to HEKn cells. These results suggest that *M. leprae* infects keratinocytes by taking advantage of the interaction of LN-5 in the basal lamina of the epidermis and a surface receptor of keratinocytes, such as α-DG, integrin-β1, or -β4.

**Author summary:** In the current study, we investigated the issue of how *M.leprae* infects keratinocytes. We focused on the role of LN-5, a predominant form of laminin of the epidermis, in the invasion of *M. leprae* in keratinocytes. Our results show that *M. leprae* preferentially binds to LN-5-coated slides and coating *M.leprae* with LN-5 enhanced its binding to human epidermal keratinocytes, neonatal (HEKn). In addition, a pre-treatment with an antibody against α-DG, integrin-β1 or -β4 inhibited the binding of LN-5-coated *M. leprae* to HEKn cells. These results suggest that *M. leprae* invades keratinocytes by taking advantage of the interaction of LN-5 in the basal lamina of the epidermis and a surface receptor of keratinocytes, such as α-DG, integrin-β1, or -β4

## Introduction

Leprosy, Hansen’s disease, is a chronic granulomatous disease caused by the intracellular bacterium *Mycobacterium leprae* (*M. leprae*). It mainly affects both the skin and peripheral nerves, resulting in the development of skin lesions, such as macules, plaques or nodules, and peripheral neuropathy [1]. *M.leprae* is usually found in macrophages and nerves of the dermal zone in patients with multibacillary leprosy [2].

In addition to the dermis, *M.leprae* can also be detected in the epidermis, sweat glands and hair follicles of patients with high bacteriological index (BI>4+) multibacillary leprosy [3]. Although leprologists generally believe that *M.leprae* is transmitted through the respiratory tract, compared to the skin route, *Job et al.* [4] reported that *M.leprae* was also present in the superficial keratin layer of the skin of lepromatous leprosy patients, suggesting that *M.leprae* may be transmitted from the intact skin of patients with lepromatous leprosy. It has been suggested that *M.leprae* is transmitted to the epidermis from rapidly growing granuloma in the upper dermis of patients with lepromatous leprosy [5]. The mechanism responsible for the epidermis invasion by *M.leprae* is not known with certainty, whereas the underlying mechanism by which *M.leprae* invades peripheral nerves, especially Schwann cells, is well defined.

*M. leprae* invades Schwann cells by binding to the alpha (α)-dystroglycan (DG) of Schwann cells via the interaction of α-DG and laminin (LN)–α2 in the basal lamina that surrounds the Schwann cell-axon unit [6]. The DG complex in Schwann cells consists of α-DG and β-DG. α-DG serves as a receptor on the Schwann cell that interacts with extracellular LN-α2, and β-DG serves as a links between the extracellular matrix (ECM) and the intracellular cytoskeleton [7, 8]. The basement membrane (BM) surrounding Schwann cells is composed of LNs, collagen IV, and proteoglycans [9]. LN-2 (α2, β1, γ1 chains) is the most common form of laminin in the basal lamina that surrounds Schwann cell-axon unit [10]. It has been reported that *M. leprae* simultaneously binds to the globular domain of LN-α2 and α-DG, a surface receptor, of Schwann cells, indicating that LN-α2 mediates the attachment and invasion of *M. leprae* to peripheral nerve cells [11].

Thus, we hypothesized that *M.leprae* uses components of the ECM, which is bound to a cell surface receptor, for the invasion of keratinocytes, as shown in Schwann cells. LN-5 (α3, β3, γ2 chains) is a major component of the basal lamina between the epidermis and dermis, and mediates the stable attachment of the epidermis to the dermis via the formation of hemidesmosomes [12]. Keratinocytes bind to LN-5, collagen, and fibronectin via integrins including α2β1, α3β1 and α6β4 [13, 14]. In addition, α/β-DG is also expressed in keratinocytes that are present in all epidermal layers except for the corneal layer [15].

In the current study, we investigated the issue of whether and how *M.leprae* invades keratinocytes. Our results show that *M.leprae* preferentially binds to LN-5 and that coating *M.leprae* with LN-5 enhanced its binding to human keratinocytes. Our results also show that a pre-treatment with antibody against α-DG, integrin-β1, or -β4 inhibited the binding of LN-5-coated *M.leprae* to human keratinocytes, suggesting that the invasion of *M. leprae* to keratinocytes is assisted by the interaction of LN-5 in the basal lamina of the epidermis and a keratinocyte surface receptor, such as α-DG, integrin-β1, or -β4.

## Materials and Methods

### Ethics statement

All procedures related to animal research were conducted in accordance with the Laboratory Animals Welfare Act, the Guide for the Care and Use of Laboratory Animals and the Guidelines and Policies for Rodent experiment provided by the IACUC (Institutional Animal Care and Use Committee) in school of medicine, The Catholic University of Korea (Approval number: CUMC-2017-0091-02). Human skin samples were obtained from patients who had upper lid blepharoplasties with no clinical evidence of inflammatory or immune diseases. These activities were undertaken after written informed consent was obtained from the donors, according to procedures approved by the Institutional Review Board of Seoul St. Mary’s Hospital (KC10TISE0743) and the tenets of the Declaration of Helsinki.

### Reagents and antibodies

Auramine O, H_2_O_2_, DAPI, Collagen IV and Fibronectin were obtained from Sigma-Aldrich (St. Louis, MO). Laminin-α2 (LN-α2, LN211-02) and laminin-5 (LN-5, ab42326) proteins were obtained from BioLamina (Matawan, NJ) and Abcam (Cambridge, MA), respectively. Antibodies against LN-5 (ab102539 for immunohistochemistry), integrin-β1 (ab24693 for immunocytochemistry and binding assay) and -β4 (ab133682 for immunocytochemistry and binding assay) were obtained from Abcam (Cambridge, MA). Antibodies against LN-α2 (sc-55605 for immunohistochemistry), α-dystroglycan (α-DG, sc-53987 for immunocytochemistry and binding assays), integrin-β2 (sc-13548 for binding assays) and –β3 (sc-52589 for binding assays) were obtained from Santa Cruz Biotechnology (Santa Cruz, CA). Cy™5-conjugated secondary antibody and horseradish peroxidase-conjugated secondary antibody were obtained from Jackson ImmunoResearch (West Grove, PA).

### *Mycobacterium leprae* (*Thai 53*) isolation

BALB/c nude mice were obtained from Orient Bio (Seong Nam, Gyunggi-do, Korea) and were maintained under specific pathogen-free conditions in the Department of Laboratory Animals, The Catholic University of Korea. Standard mouse chow (Ralston Purina, St Louis, MO) and water were provided ad libitum. The foot-pads of *M. leprae*-infected BALB/c *nude* mice were treated with potadine solution and washed with ice-cold Dulbecco’s phosphate-buffered saline (DPBS, Sigma-Aldrich Co. Ltd, MO) to remove exogenous contamination. The foot-pads were excised, cut into small pieces, and homogenized with a MACs isolator (Miltenyl Biotec, Teterow, Germany). The extract was filtered using a cell strainer (BD Falcon, Durham, NC) to remove tissue debris and centrifuged at 3,000 rpm (Rotanta 460R, Hettich, Japan) for 25 min at 4 °C. The pellet was resuspended in 1 ml of ice-cold DPBS and treated with 2 N sodium hydroxide for 5 min. The reaction mixture was neutralized by adding 13 ml of ice-cold DPBS (Sigma-Aldrich Co. Ltd, MO). After centrifugation and resuspension, acid-fast bacillus (AFB) staining was performed and the number of bacteria counted by light microscopy under an oil immersion field using a procedure established by Shepard and McRae.

### Cell cultures

Human primary epidermal keratinocytes from neonatal foreskin (HEKn) cells were acquired from Invitrogen (Carlsbad, CA) and grown in EpiLife medium supplemented with 100 U/ml of penicillin, 100 mg/ml of streptomycin, 250 ng/ml of amphotericin B, 60 μM of calcium, and Human keratinocyte growth supplement (HKGS, Cascade Biologics; Invitrogen, Carlsbad, CA). These cells were maintained in a state of proliferation and non-differentiation. The cells were passaged with a gentle TrypLE select (Invitrogen, Carlsbad, CA) treatment followed by a trypsin neutralization solution (Invitrogen, Carlsbad, CA). The cells were plated on 6-well plates at 1 × 10^5^ cells/well or onto 4-channel chamber slides (Lab-Tek II chamber slide, Thermo Fisher Scientific, Waltham, MA) at 5 × 10^4^ cells/well and were grown until reaching 70% confluence in serum-free EpiLife medium supplemented with HKGS.

### Infection of HEKn cells with *M. leprae*

The HEKn cells were cultured on coverslide in a 6-well plate. *M. leprae* was pre-incubated with LN-α2 (10 μg/ml) or LN-5 (2 μg/ml) in DPBS for 2 h at 37 °C, followed by washing. The cells were infected with *M. leprae* at multiplicities of infection (MOI) of 10:1, 20:1, 50:1 and 100:1 for 1 h at 37 °C. After removing extracellular *M. leprae* by washing with phosphate-buffered saline (PBS), *M. leprae* were stained with the AFB stain or Auramine O, and examined in an oil immersion field of a light microscopy or fluorescence microscopy.

### Immunohistochemistry

The skins were fixed in 4% formaldehyde for 4 h at room temperature prior to embedding in paraffin and 4 μm thick sections were dewaxed and rehydrated in a graded series of alcohol solutions. The sections were incubated in 0.3% sodium citrate buffer (pH 6.0) for 10 min at 100 °C and 3% hydrogen peroxide (H_2_O_2_) for 10 min after which, they were rinsed with PBS and incubated in blocking solution [5% goat serum and 0.001% Tween-20 in tris-buffered saline (TBS)] for 20 min. The sections were then incubated overnight with an antibody against LN-α2 or LN-5 in an incubation solution (5% goat serum and 0.1% Tween-20 in TBS) at 4 °C. After washing with PBS, the sections were incubated with a mouse Cy™5- or a rabbit Cy™5-conjugated secondary antibody at room temperature for 2 h. Nuclei were counterstained for 5 min with DAPI (Sigma-Aldrich Co. Ltd, MO). The negative control was processed in the absence of the primary antibody. Immunofluorescence was visualized by confocal microscopy (LSM 510 Meta, Zeiss, Germany).

### Immunocytochemistry

The cells were fixed in 4% paraformaldehyde in PBS. The fixed cells were then rinsed with PBS and incubated in blocking solution (5% goat serum and 0.001% Tween-20 in TBS) for 20 min. The cells were then incubated overnight with an antibody against α-DG, integrin-β1, or -β4 in an incubation solution (5% goat serum and 0.1% Tween-20 in TBS) at 4 °C. After washing with PBS, the cells were incubated with a mouse Cy™5- or a rabbit Cy™5-conjugated secondary antibody at room temperature for 2 h. Nuclei were counterstained for 5 min with DAPI (Sigma-Aldrich Co. Ltd). The negative control was processed in the absence of the primary antibody. Immunofluorescence was visualized by confocal microscopy (LSM 500 Meta, Zeiss, Germany).

### Bacterial adherence assays

In the assay for the binding of *M.leprae* to the ECM-coated culture plate, 4-channel chamber slides were coated, as described in a previous report [16]. The slides were coated with 0.1 μg/ml of LNs, type IV collagen or fibronectin by incubation at room temperature overnight. Saline was used as a negative control. Nonspecific binding was blocked with 5% BSA for 3 h at 37 °C and the sample then washed 5 times with DPBS. Ten microliters of a suspension of *M.leprae* (5 × 10^8^ bacteria/ml) was added to each well followed by incubation for 1 h at 37 °C. Unbound bacteria were removed by washing 5 times with DPBS. After fixation with 2% paraformaldehyde for 10 min, the bacteria were stained with Auramine O. The level of Auramine O-labeled *M. leprae* that was bound to slide was determined using the ZEN program (Zeiss, Oberkochen, Germany) under a LSM 510 Meta confocal microscopy (Zeiss, Oberkochen, Germany).

For assaying the binding of *M.leprae* to HEKn cells, the HEKn cells were cultured in 4-channel chamber slides and incubated overnight at 37 °C under 5% CO_2_. For determining the *M. leprae* that was bound to HEKn cells, *M. leprae* was pre-incubated with 10 μg/ml LN-α2 or 2 μg/ml LN-5 for 2 h at 37 °C before inoculation at MOI of 10:1, 20:1, 50:1 and 100:1. For the binding inhibition assay, HEKn cells were pre-incubated with an antibody against α-DG, integrin-β1, -β2, -β3 or –β4 for 2 h at 37 °C before inoculation with *M. leprae* at an MOI of 100:1. After inoculating the HEKn cells with *M. leprae* for 1 h at 37 °C in 5% CO_2_, extracellular *M. leprae* were removed by washing 5 times with PBS and fixing in 2% paraformaldehyde for 30 min. *M. leprae* were labeled with the AFB stain and examined in the oil immersion field of a light microscopy.

## Results

### HEKn cells were infected with *M. leprae*

We initially investigated the issue of whether *M.leprae* infects HEKn cells. HEKn cells were incubated with *M. leprae* at multiplicity of infections (MOI) of 10:1, 20:1, 50:1 and 100:1, respectively, for 6 h at 37 °C. At an MOI of 100:1, 77.4% of the cells were infected with *M. leprae* and the average number of *M.leprae* per cell was 3 (Fig 1).

**Fig 1.**
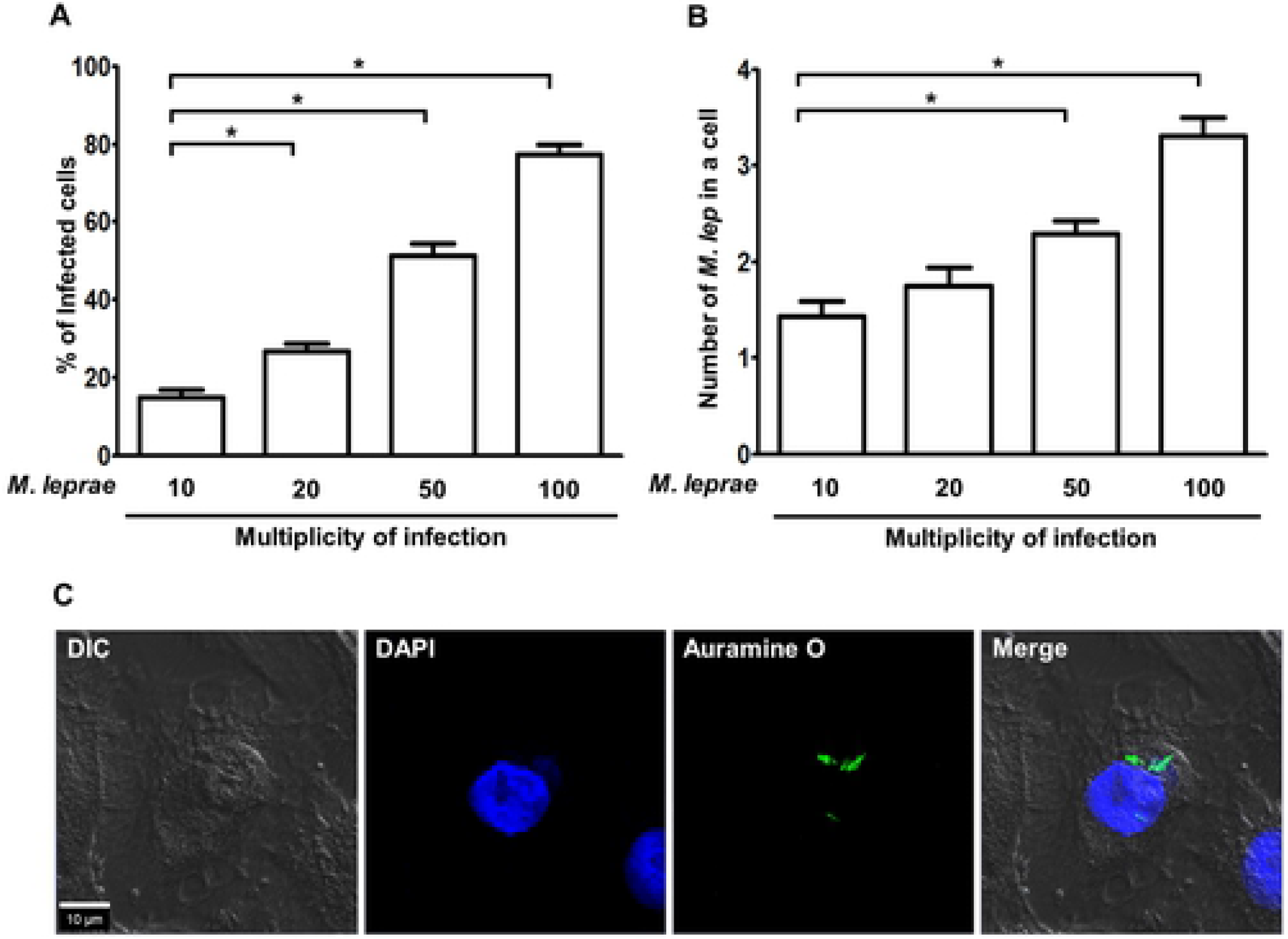
HEKn cells were infected with *M.leprae*. (A and B) HEKn cells were incubated with *M. leprae* at MOI of 10:1, 20:1, 50:1 and 100:1, respectively, for 6 h at 37°C. After removing extracellular *M. leprae* by washing, the sample was stained with AFB stain. The percentage of *M. leprae* that infected the cells and the number of *M. leprae* per a cell were determined in the oil immersion field of a light microscopy. Significance was calculated by a one way ANOVA and Tukey’s multiple comparison tests. \**P <0.05* versus cells were incubated with *M. leprae* at the MOI of 10:1. (C) HEKn cells were incubated with *M. leprae* at MOI of 100:1 for 6 h at 37 °C. After removing extracellular *M. leprae* by washing, the preparation was stained with Auramine O. Nuclei were counterstained for 5 min with DAPI. Scale bar: 10 μm.

### LN-5, but not LN-α2, was expressed in the basal lamina of the human epidermis and α-DG, integrin-β1 and -β4 were expressed in HEKn cells

We examined the expression pattern of LN-α2 and LN-5 in human skin. Consistent with previous reports [12], LN-5, but not LN-α2, was expressed in the basal lamina between the epidermis and dermis (Fig 2). We then examined the expression patterns of cell surface receptors in HEKn cells. As shown in Fig 3, HEKn cells expressed α-DG, integrin-β1 and -β4 on the cell surface.

**Fig 2.**
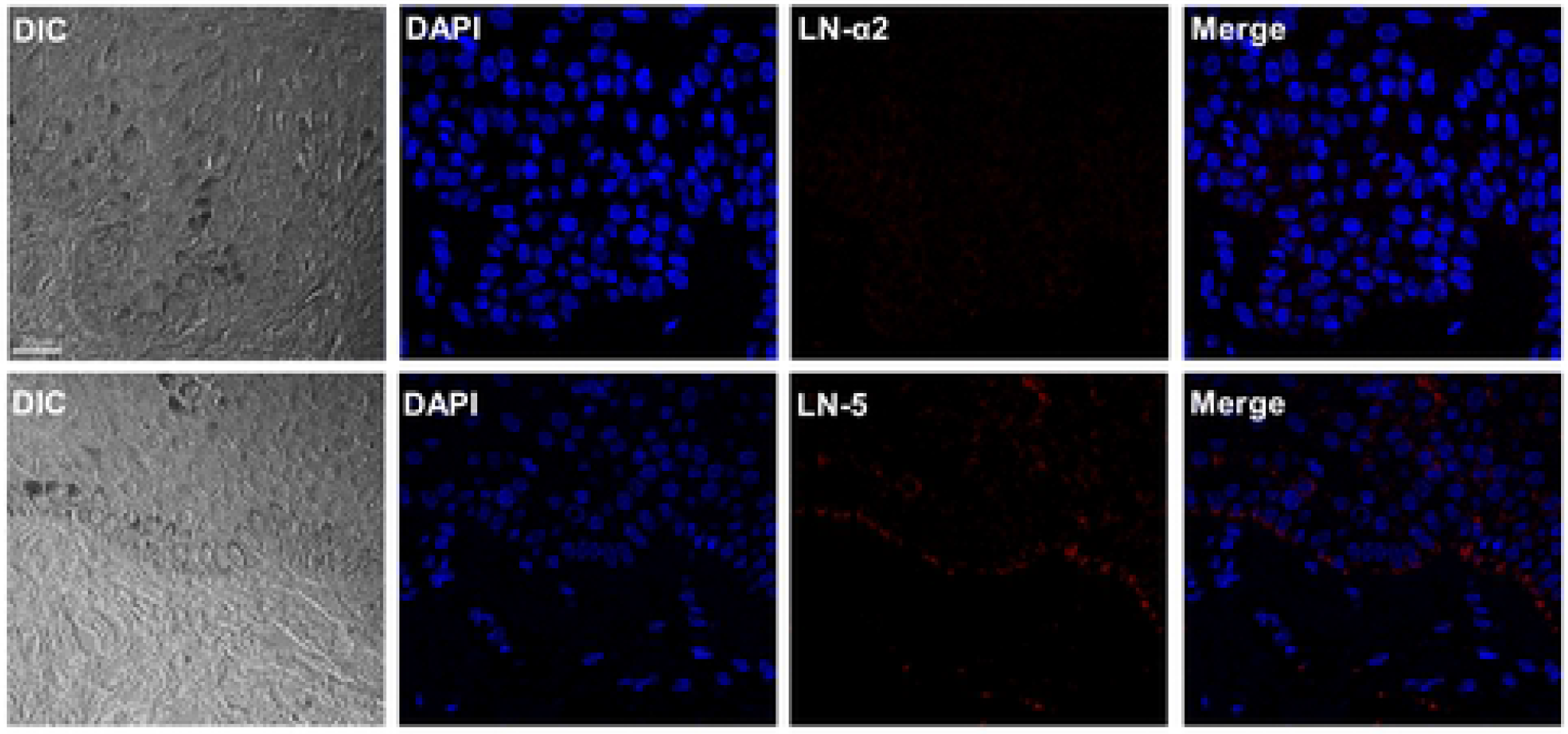
LN-5, but not LN-α2, was expressed in human epidermis. Human skin was immunostained with an antibody against LN-α2 or LN-5. After washing with PBS, the skin samples were incubated with a mouse Cy™5- or a rabbit Cy™5-conjugated secondary antibody at room temperature for 2 h. Nuclei were counterstained for 5 min with DAPI. Scale bar: 20 μm.

**Fig 3.**
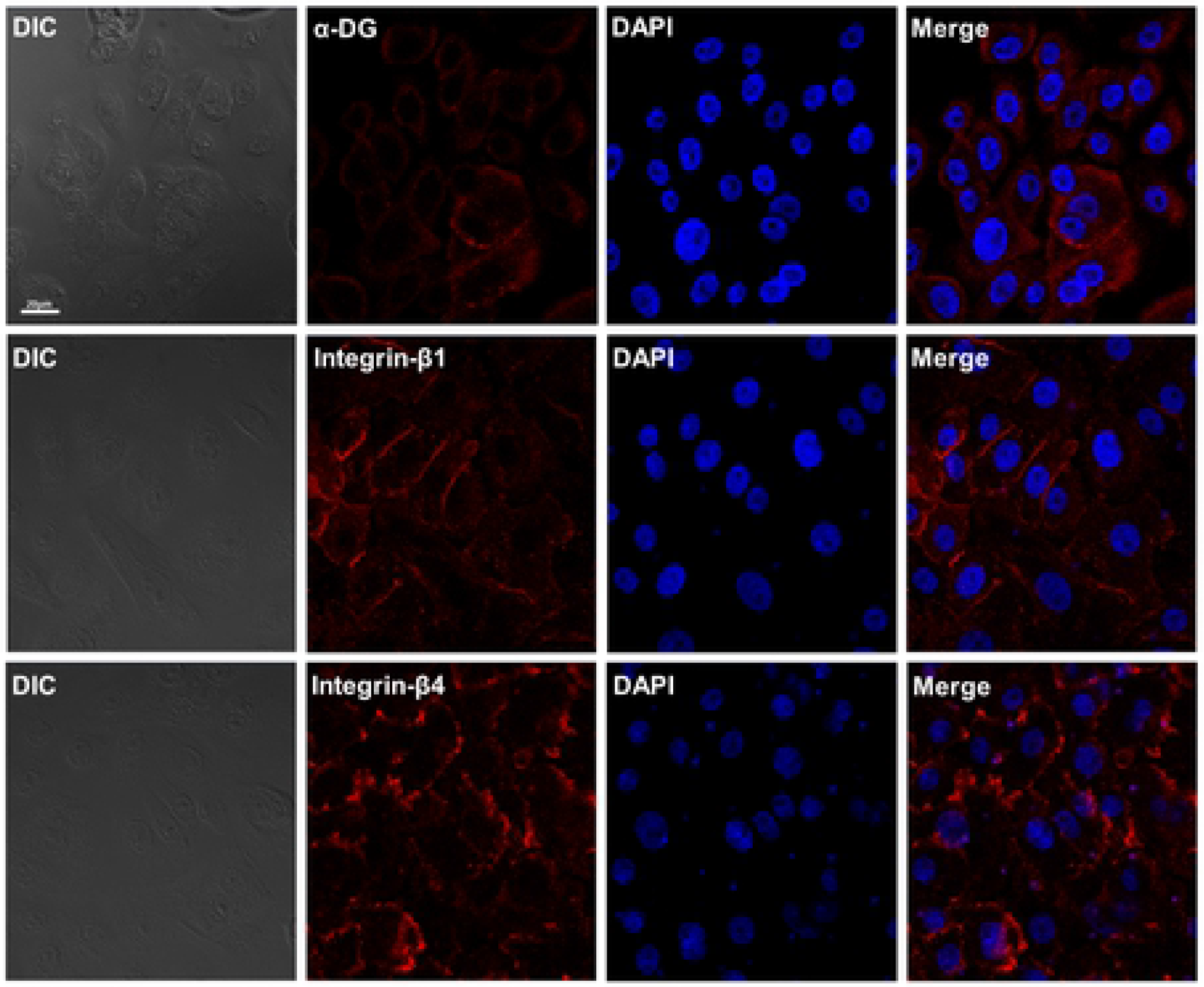
α-DG, integrin- β1 and -β4 were expressed in HEKn cells. HEKn cells were immunostained with an antibody against α-DG, integrin-β1 or -β4, respectively. After washing with PBS, the HEKn cells were incubated with a mouse Cy™5- or a rabbit Cy™5-conjugated secondary antibody at room temperature for 2 h. Nuclei were counterstained for 5 min with DAPI. Scale bar: 20 μm.

### Coating of *M. leprae* with LN-5 enhanced the binding of *M. leprae* to HEKn cells

We then investigated the issue of whether *M. leprae* adheres to the immobilized extracellular matrix LN-5, collagen IV and fibronectin using a solid-phase bacterial-adherence assay. We used LN-α2 as a positive control since LN-α2 in Schwann cells basal lamina is known to be the primary target molecule for *M. leprae* [16].

The level of *M. leprae* binding was increased in the LN-α2- as well as the LN-5-coated slides, compared to collagen IV- and fibronectin-coated slides (Fig 4). We also examined the binding ability of LN-α2- or LN-5-coated *M.leprae* to HEKn cells. As shown in Fig 5, the coating of *M. leprae* with LN-α2 or LN-5 resulted in an increase in the number of *M. leprae* that had adhered HEKn cells (average number of adherent *M. leprae* to HEKn cells per 100 HEKn cells; 69.3±5.7 in LN-α2-coated *M. leprae* and 44.0±2.4 in LN-5-coated *M. leprae* in comparison with 35.0±3.2 in non-treated *M. leprae*).

**Fig 4.**
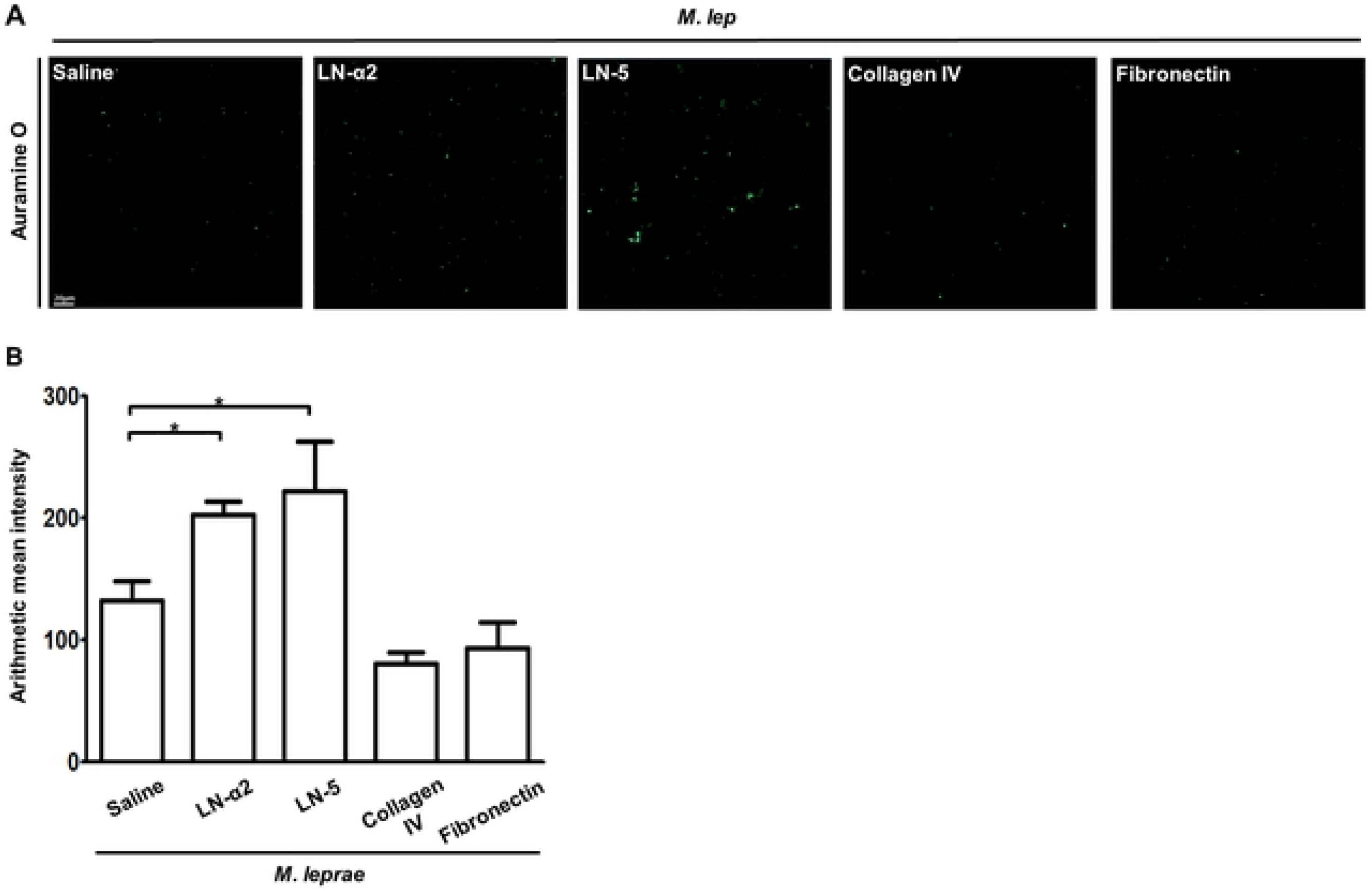
*M. leprae* preferentially bound to LN-α2 and LN-5, compared to collagen IV and fibronectin. (A and B) *M. leprae* (5 × 10^8^) were overlaid onto LN-α2, LN-5, collagen IV, or fibronectin-coated 4-chamber slides and incubated for 1 h at 37 °C. After removing unattached *M. leprae* by washing, the *M. leprae* were stained with Auramine O. The level of binding activity of *M. leprae* to ECM-coated slides was determined by measuring Auramine O fluorescence activity. \**P <0.05* between the indicated groups. Scale bar: 20 μm.

**Fig 5.**
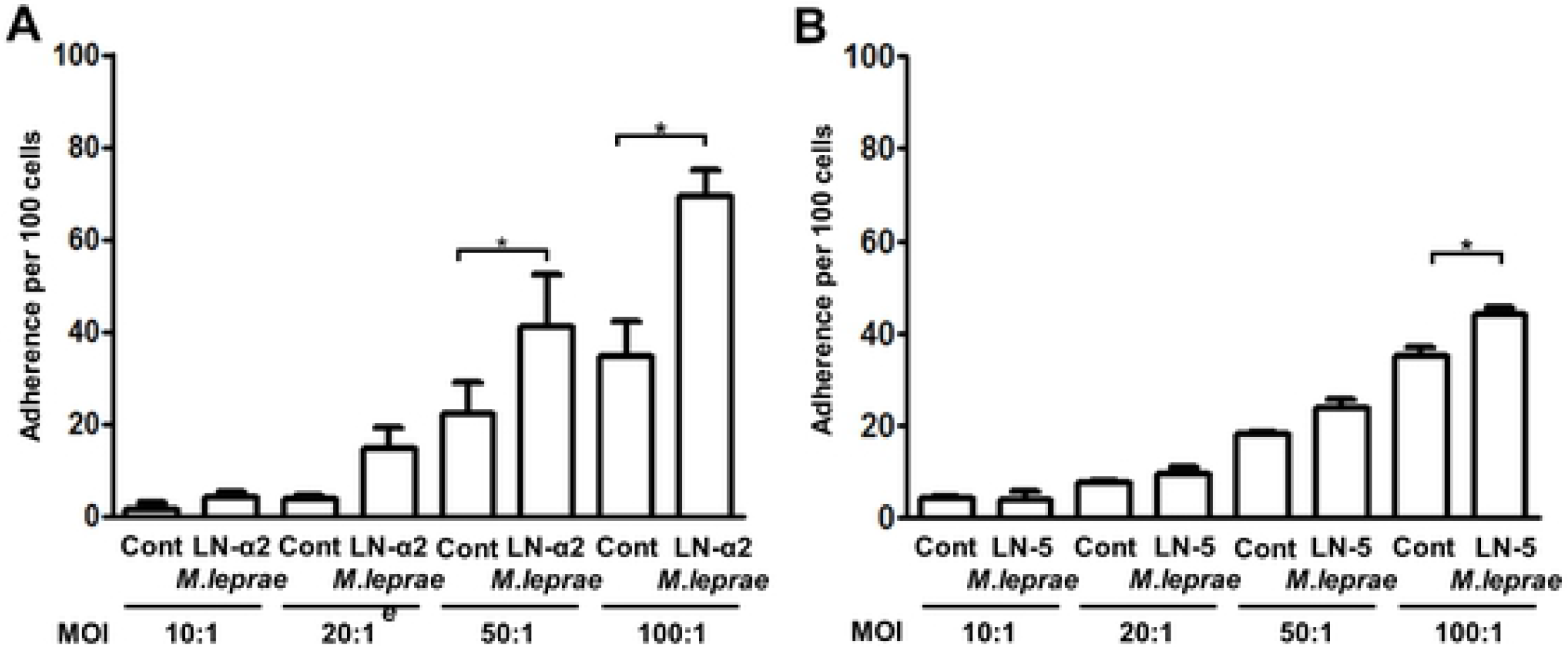
Coating *M. leprae* with LN-α2 or LN-5 enhanced the binding of *M. leprae* to HEKn cells. (A and B) HEKn cells were incubated with LN-α2- (A) or LN-5- (B) coated *M. leprae* at MOI of 10:1, 20:1, 50:1 and 100:1, respectively, for 1 h at 37 °C. After removing unattached *M. leprae* by washing, the samples were stained with the AFB stain. The number of *M. leprae*–bound cells per 100 HEKn cells was determined in the oil immersion field of light microscopy. \**P <0.05* between the indicated groups.

### Pre-treatment with antibody against α-DG, integrin-β1, or -β4, inhibited binding of LN-5-coated *M. leprae* to HEKn cells

Rambukkana et al. [6] reported that when *M. lerpae*, that had been coated with the recombinant globular domain of LN-α2 (LN-α2G), were pre-incubated with recombinant α-DG, the LN-α2G/α-DG-mediated *M.leprae* binding to rat Schwann cells was competitively inhibited, suggesting the existence of a linkage between LN-α2 and α-DG in the interaction of *M.leprae* with Schwann cells. In the current study, although LN-α2 is not expressed in skin, we employed the LN-α2/α-DG-mediated *M.leprae* binding to cells as a positive control in the binding assay. Consistent with Rambukkana et al.’s results [6], our result also showed that the pre-treatment of HEKn cells with an anti-α-DG antibody inhibited the binding of LN-α2-coated *M.leprae* to HEKn cells (Fig 6A). In addition, the pre-treatment of HEKn cells with antibody against α-DG, integrin-β1, or -β4, all of which are expressed on the surface of HEKn cells (Fig 3), inhibited LN-5-coated *M. leprae* from binding to HEKn cells (Fig 6B and C). However, pre-treatment with antibody against integrin-β2 or -β3 had no effect on inhibiting the binding of LN-5-coated *M.leprae* to HEKn cells (Fig 6B). These results suggest that *M. leprae* invades keratinocytes by taking advantage of the interaction of LN-5 in the basal lamina of the epidermis and a surface receptor of keratinocytes, such as α-DG, integrin-β1, or -β4.

**Fig 6.**
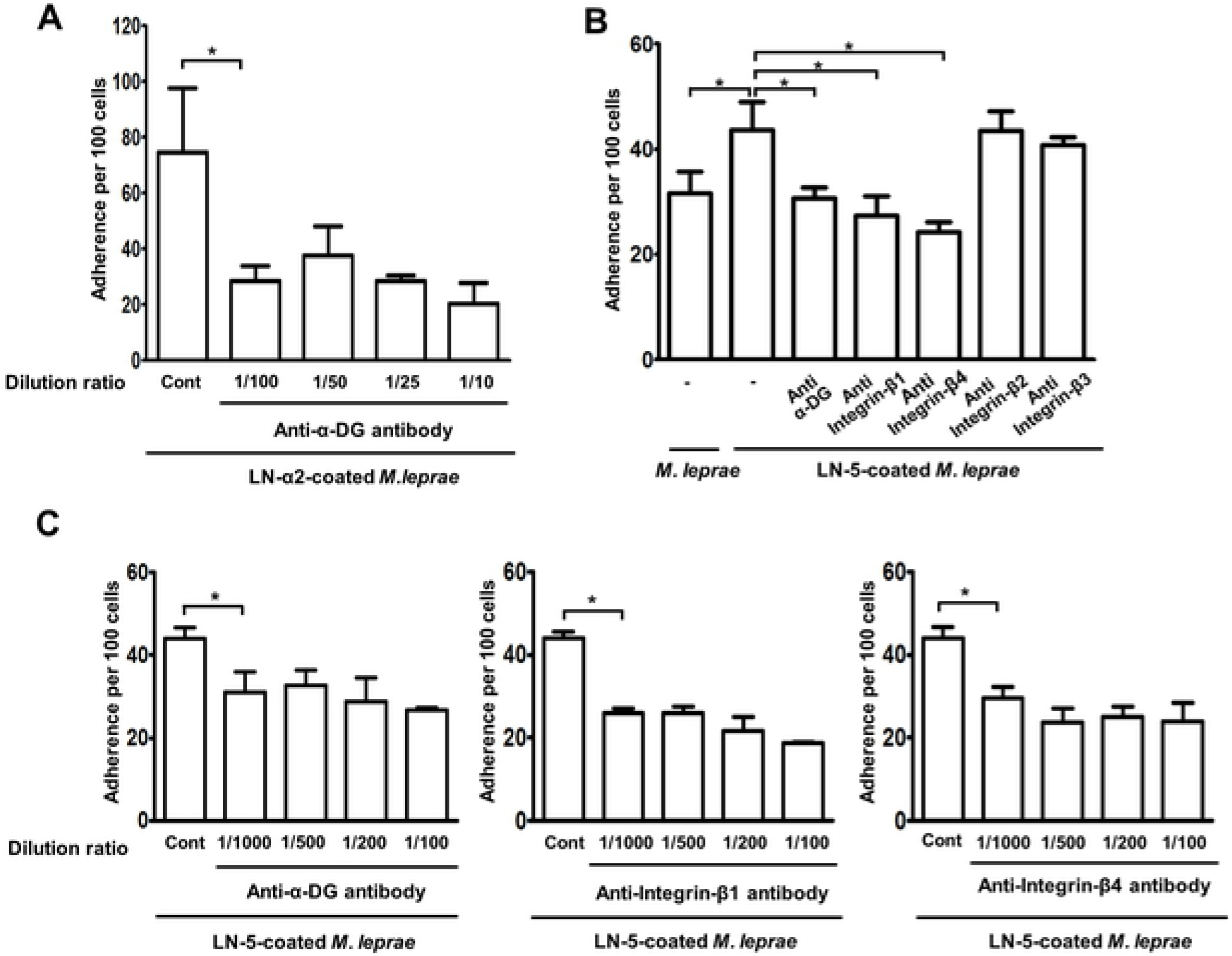
Pre-treatment with an antibody against α-DG, integrin-β1 or -β4 inhibited binding of LN-5-coated *M. leprae* to HEKn cells. (A) HEKn cells were pre-treated with an antibody against α-DG at the indicated dilution ratio for 1 h, and then further incubated with LN-α2-coated *M. leprae* at MOI of 100:1 for 1 h at 37 °C. (B and C) HEKn cells were pre-treated with an antibody against α-DG, integrin-β1, -β2, -β3, or -β4, at the dilution ratio 1:1,000 (B) and at the indicated dilution ratio (C) for 1 h. HEKn cells were then incubated with LN-5-coated *M. leprae* at MOI of 100:1 for 1 h at 37 °C. After unattached *M. leprae* were washed away, *M. leprae* were stained with the AFB stain. The number of *M. leprae*–bound cells per 100 HEKn cells was determined in the oil immersion field of light microscopy. \**P <0.05* between the indicated groups.

## Discussion

ECM is an acellular proteinaceous fraction of the tissues. ECM proteins consist of collagen, elastin, fibrillin, LNs, fibronectin, vitronectin, thrombospondin, proteoglycans and hyaluronic acid. ECM is involved in the structural support of tissues as well as various cellular signaling processes, including cell adhesion, migration, growth, and differentiation [17]. Although pathogens need to breach and degrade ECM proteins in order to successfully invade a tissue, they also utilize ECM proteins to aid in their adhesion to host tissues. LNs and collagens are major target glycoproteins of various pathogens, such as bacteria, fungi, and viruses, for adhesion to cells of host tissue [18].

LNs are heterotrimeric glycoproteins that consist of α, β and γ chain. The chains, α, β and γ, which are connected to one another via disulfide bonds at their C-terminal regions, form a triple coiled-coil region, resulting in a ‘crucifix’-shaped structure [18]. There are currently five α chain, three β chain and three γ chain isoforms and 16 LN isoforms have been identified in humans [19]. LN isoforms are differentially distributed in human tissues or cells [18]. LN-2 (α2, β1, γ1 chains) is a predominant laminin associated with Schwann cells [10]; LN-5 (α3, β3, γ2 chains) is found in oral, intestinal and dermal epithelial cells [12, 20, 21]; LN-10/11 is expressed in the lung epithelium [22]. The interaction between ECM laminins and integrins of epithelial cells confers mechanical stability to tissues as well as an invasive mechanism for pathogens [18].

It has been reported that *M. leprae* binds to the globular domains (LG1, LG4, and LG5 domains) of LN-α2 chain and that the LN-α2 chain simultaneously binds to α-DG, a surface receptor, of Schwann cells, resulting in the attachment and invasion of *M. leprae* to Schwann cells [11]. Our results also show that coating *M. leprae* with LN-α2 enhanced the binding of *M. leprae* to HEKn cells (Fig 5A) and a pre-treatment with an antibody against α-DG inhibited the binding of LN-α2-coated *M. leprae* to HEKn cells (Fig 6A). However, although LN-α2 (α2, β1, γ1 chains) mediates the attachment of *M. leprae* to HEKn cells, it was not detected in the skin (Fig 2), whereas LN-5 (α3, β3, γ2 chains) is a major form of laminins that is present between the epidermis and dermis [12]. Thus, in the current study, we focused on the role of LN-5 in the invasion of *M. leprae* to keratinocytes.

It has been reported that LN-5, which is expressed in the BM between the epidermis and dermis, has been reported to be a target molecule and mediator for the invasion of the Human papilloma virus (HPV) to keratinocytes [23]. HPV first infects keratinocytes in the basal layer of the epithelium and then replicates in a fully differentiating squamous epithelium [24]. Culp et al. [23] reported that the HPV capsid binds to LN-5 in the ECM of culture keratinocytes. In that report, the authors reported that, when sections of cervical mucosa tissues were incubated with HPV, the HPV became bound to the suprabasal layer and BM of the cervical mucosa and that a pre-treatment with anti-LN-5 antibody blocked the binding of HPV to these cervical mucosa tissue sections. Our results also show that *M. leprae* preferentially bound to LN-5-coated slides, compared to collagen IV and fibronectin (Fig 4) and that coating *M. leprae* with LN-5 enhanced the binding of *M. leprae* to HEKn cells (Fig 5B), suggesting LN-5 mediates the attachment and invasion of *M. leprae* to HEKn cells. Although *M.leprae* can be detected in the all layers of the skin, it is more frequently detected in the suprabasal and basal layers of the epidermis of patients with multibacillary leprosy [3, 25]. We conclude that the clinical findings support the conclusion that LN-5 in the BM of the epidermis mediates the attachment and invasion of *M.leprae* to non-differentiated, proliferating keratinocytes in the basal layer. In the current study, to limit the differentiation of HEKn cells, we maintained HEKn cells in EpiLife medium supplemented with human keratinocyte growth supplement (HKGS, Cascade Biologics; Invitrogen, Carlsbad, CA), and not in fetal bovine serum.

It is well known that α-DG serves as a Schwann cell receptor for the LN-α2-mediated *M.leprae* invasion of Schwann cells [6]. In the skin, DG is present in the epidermal BM [26]. Thus, we hypothesized that α-DG is also involved in the LN-5-mediated *M.leprae* invasion of keratinocytes, as shown in the LN-2α-mediated *M.leprae* invasion of Schwann cells. As shown in Fig 6B, our results show that pre-treatment with an anti-α-DG antibody blocked the binding of LN-5-coated *M.leprae* to HEKn cells. LN-5 permits the stable attachment of the epidermis to the dermis via interaction with α/β-DG as well as integrins of keratinocytes [12–14, 26]. In addition, the interaction of LN-5 with integrin α3β1 and α6β4 activates the adhesion and spreading of keratinocytes for wound healing [13, 14]. These previous results indicate that LN-5/α3β1 or α6β4 may be involved in mediating the attachment of *M.leprae* to HEKn cells and their subsequent invasion. Consistent with these results, the findings reported herein show that a pre-treatment with anti-integrin β1 or β4 antibody blocked the binding of LN-5-coated *M.leprae* to HEKn cells (Fig 6B).

Although *M.leprae* is not frequently detected in the epidermis, studies have clearly shown that *M.leprae* is found in the epidermis of patients with multibacillary leprosy [3, 4, 25, 27–29]. *M.leprae* was detected in all layers of the epidermis, basal, suprabasal, prickle cells, and keratin layers [3, 4]. In addition, *M.leprae* was also reported to be distributed in sweat glands and hair follicles [3]. Job et al. [4] suggested that the transepidermal discharge of *M.leprae* may be attributed to the possibility that *M.leprae* is transferred to the keratin layer by travelling inside keratinocytes from the basal to the keratin layer and that *M.leprae* then exits from hair follicles or sebaceous glands. Satapathy et al. [25] suggested that health workers in leprosy control should consider the possibility that leprosy can be transmitted through the skin and by skin to skin contact, since large numbers of *M.leprae* are shed, even through intact skin.

The findings reported in this study suggest that *M. leprae* invades non-differentiated, proliferating HEKn cells by taking advantage of the interaction of LN-5 in the basal lamina of the epidermis and a surface receptor on keratinocytes, such as α-DG, integrin-β1, or -β4.

## Acknowledgments

We wish to thank Park Jieun (Integrative Research Support Center, The Catholic University of Korea) for the immunofluorescence analysis. This research was supported by grants from Korea Centers for Disease Control & Prevention, Korea Ministry of Health & Welfare (5-2018-A0082-00005).

